# Automated classification of synaptic vesicles in electron tomograms of *C. elegans* using machine learning

**DOI:** 10.1101/291310

**Authors:** Kristin Verena Kaltdorf, Maria Theiss, Sebastian Matthias Markert, Mei Zhen, Thomas Dandekar, Christian Stigloher, Philip Kollmannsberger

## Abstract

**Abstract:** Synaptic vesicles (SVs) are a key component of neuronal signaling and fulfil different roles depending on their composition. In electron micrograms of neurites, two types of vesicles can be distinguished by morphological criteria, the classical “clear core” vesicles (CCV) and the typically larger “dense core” vesicles (DCV), with differences in electron density due to their diverse cargos. Compared to CCVs, the precise function of DCVs is less defined. DCVs are known to store neuropeptides, which function as neuronal messengers and modulators [1]. In *C. elegans*, they play a role in locomotion, dauer formation, egg-laying, and mechano- and chemosensation [2]. Another type of DCVs, also referred to as granulated vesicles, are known to transport Bassoon, Piccolo and further constituents of the presynaptic density in the center of the active zone (AZ), and therefore are important for synaptogenesis [3].

To better understand the role of different types of SVs, we present here a new automated approach to classify vesicles. We combine machine learning with an extension of our previously developed vesicle segmentation workflow, the ImageJ macro 3D ART VeSElecT. With that we reliably distinguish CCVs and DCVs in electron tomograms of *C. elegans* NMJs using image-based features. Analysis of the underlying ground truth data shows an increased fraction of DCVs as well as a higher mean distance between DCVs and AZs in dauer larvae compared to young adult hermaphrodites. Our machine learning based tools are adaptable and can be applied to study properties of different synaptic vesicle pools in electron tomograms of diverse model organisms.

**Author summary:** Vesicles are important components of the cell, and synaptic vesicles are central for neuronal signaling. Two types of synaptic vesicles can be distinguished by electron microscopy: the classical “clear core” vesicles (CCVs) and the typically larger “dense core” vesicles (DCVs). The distinct appearance of vesicles is caused by their different cargos. To rapidly distinguish between both vesicle types, we present here a new automated approach to classify vesicles in electron tomograms. We combine machine learning with an extension of our previously developed vesicle segmentation workflow, an ImageJ macro, to reliably distinguish CCVs and DCVs using specific image-based features. The approach was trained and validated using data-sets that were hand curated by microscopy experts. Our technique can be transferred to more extensive comparisons in both stages as well as to other neurobiology questions regarding synaptic vesicles.

## 3. Introduction

*Caenorhabditis elegans* is a well-studied model organism. Its small nervous system, consisting of 302 neurons [4,5] allows for studies of the connectome in its entirety [5–7]. A focus of research is on *C. elegans* young adult hermaphrodites, which makes them a reference system for comparison with other developmental states in the life cycle. The most remarkable of all *C. elegans* developmental states is the dauer larva.

When the L1 stage larvae sense adverse environmental conditions such as heat, overpopulation and starvation, they activate an alternative state, diapause, instead of continuing the reproductive life cycle [8,9]. Dauer larvae are characterized by their ability to survive several months without any food uptake [10]. Although metabolic activity is reduced [10], dauer larvae are able to move quickly, if necessary, in particular to escape adverse environmental stimuli and in order to reach and colonize new habitats. Improvement of environmental conditions leads to dauers to exit diapause, resume development into pre-adult L4 larvae, and re-enter the reproductive life cycle [9]. The mechanisms behind this strikingly fast developmental transition, and the role of the dauer nervous system in sensing and responding to environmental cues, remain largely enigmatic.

Neuronal signaling plays a critical role in regulating the dauer stage. Neuronally released insulin-like peptides [11–14] and morphogens such as TGF-beta [15,16] regulate the activation of dauer diapause. These observations suggest potential changes of the nature of neuronal signaling in dauer larvae.

Neuropeptides are stored and released through dense core vesicles (DCVs) [1,17,18]. Recent studies, particularly the electron microscopic reconstruction of *C. elegans* prepared by high-pressure freezing, reveal that almost all synapses, including the neuromuscular junctions (NMJs), contain two principal types of vesicles, a small fraction of DCVs and the much more abundant clear core vesicles (CCVs) [19–21]. CCVs are characterized by their clear inner core in electron micrographs, which stands in strong contrast to the stained vesicle membrane. CCVs typically contain neurotransmitters such as acetylcholine and GABA in the *C. elegans* NMJs that are released into the synaptic cleft after fusion of the CCV with the presynaptic cell membrane at the release sites at active zones [2]. In contrast to CCVs, the role of DCVs is much more enigmatic. DCVs are characterized by their eponymous dark, electron dense core that typically contains neuropeptides, small protein molecules with about 3 - 100 amino acids [1] or other protein cargos such as active zone proteins [3]. DCVs have a bigger average size of about 60 - 80 nm in comparison to CCVs, which have a much smaller diameter [19], that can vary in different organisms or developmental states [22,23]. Whereas CCVs cluster around the active zone, DCVs tend to locate further away from the release site, typically at the rim of the SV pool as described for *C. elegans* NMJs [19].

DCV exocytosis is considered to mediate slow synaptic and non-synaptic signaling in neurons [1]. It typically occurs independently of the active zone and only at high rates of neuronal activity [1,2]. In *C. elegans*, neuropeptides participate in a wide variety of behaviors such as egg-laying, locomotion, dauer formation, mechano- and chemosensation depending on the expression pattern of the peptides [2]. Despite the known critical involvement of neural peptides in dauer development, whether the dauer states exhibit differences in DCV distribution in the nervous system remains unknown.

We used high-resolution 3D transmission electron microscopy (electron tomography) to systematically investigate differences of DCV and CCV in NMJs of *C. elegans* young adult hermaphrodites and dauer larvae. Series of 2D images were taken from different viewing angles by tilting the sample, and reconstructed *in silico* to obtain one 3D image stack with near isotropic resolution of typically 4 – 5 nm, determined by measuring the “unit membrane” thickness [19,25–28]. Quantification of SV in 3D electron tomograms requires reconstruction of the vesicle pool of a sufficient number of tomograms, which is very time-consuming and prone to mistakes when performed manually.

To circumvent these shortcomings we previously developed *3D ART VeSElecT*, a workflow for automated reconstruction of vesicles [22]: This ImageJ/Fiji tool allows for automated reconstruction of synaptic vesicle pools in electron tomograms, but is not able to differentiate between different types of vesicles. To solve this issue, we here present a python-based machine learning approach that trains vesicle classifiers, and a straightforward ImageJ/Fiji tool that applies a trained classifier to previously detected vesicles.

We show that our technique can differentiate between DCV and CCV in two developmental states, young adult hermaphrodites and dauer larvae. The application of these image analysis and machine learning tools for electron tomograms is in principle useful to study differences in synaptic vesicle pools of many other model organisms.

## 4. Results

### 4.1 Manual classification shows three distinct vesicle types

To compare vesicle properties in two different developmental states of *C. elegans*, dauer larvae and young adult hermaphrodites, we used high-pressure freezing and subsequent freeze substitution to immobilize and fix animals in a near-to-native state. Using electron tomography on ~250 nm cross-sections of the worm, we obtained high resolution 3D images that allow for investigation of the ultrastructure of the nematode’s NMJs. We used this information to compare synaptic vesicle pool characteristics, aiming for an automated approach for vesicle classification.

Upon visual examination of electron microscopic datasets of the NMJs of *C. elegans* dauer samples, we noted an increased fraction of DCV in dauer larvae compared to young adult hermaphrodites. To systematically quantify this, vesicles were automatically extracted from tomograms, using an improved version of our 3D ART VeSElecT workflow (see S1 Fig), and then manually assigned for vesicle types by two experts. In uncertain cases, a third expert was consulted. Since closer examination of tomograms revealed that a definite assignment of some vesicles to one of the two classes (CCV and DCV) was difficult, we initially established a third, hypothetic group of non-determinable (ND) vesicles. Non-assignable (NA) vesicles are a subgroup of ND that we could ultimately not classify as CCV or DCV. Fig 1 shows a representative selection of clear core, non-determinable and dense core vesicles from 8 exemplary tomograms (4 dauer larva tomograms and 4 hermaphrodite tomograms).

**Fig 1.**
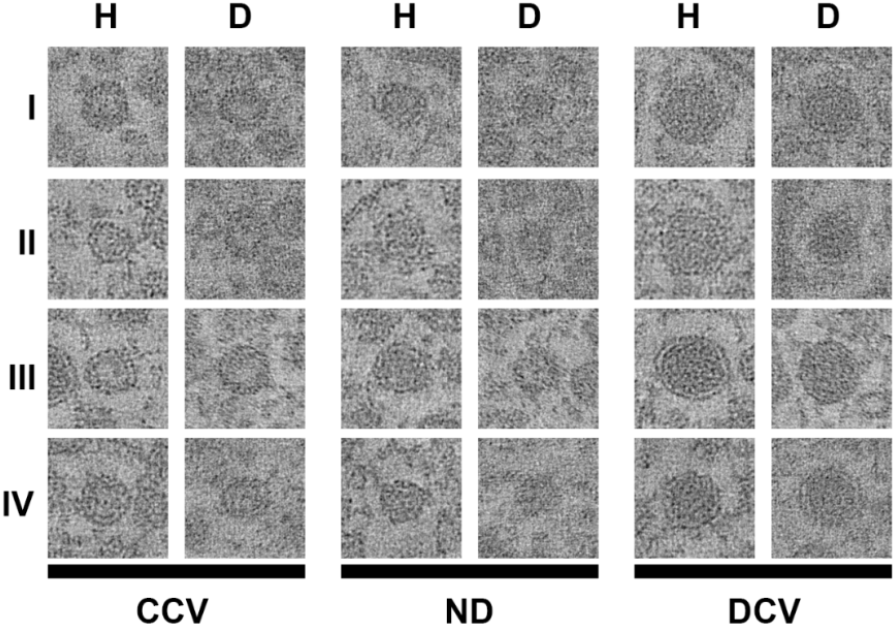
Comparison of CCV, ND vesicles and DCV of dauer larvae and young adult hermaphrodites. A selection of representative pictures is shown. Pictures of the same developmental state (young adult hermaphrodite (H) / dauer larva (D)) and with identical number (I – IV) are from the same tomogram (e.g. CCV DI, ND DI and DCV DI originate from exactly the same tomogram). Edge length of micrographs = 100 nm.

Together, we included 1072 segmented vesicles from the 7 young adult hermaphrodite tomograms, and 1108 segmented vesicles from the 8 dauer larva tomograms for analyses (Table 1). When we included all identified vesicle populations from all tomograms, the percentage of manually labeled DCVs differed slightly for dauer larvae (13%) and adult hermaphrodite (10%) datasets. After filtering the datasets, comparing only the vesicle population from the same type of NMJ with the bouton fully reconstituted in the tomograms (putative cholinergic NMJs with central cut through the AZ area and at least 100 vesicles), the dauer larvae NMJs exhibited a higher DCV ratio of 16% versus 9% in young adult hermaphrodites.

**Table 1:** number and percentage of vesicle types in all tomograms^1^

| Developmental state | Tomograms (n) | Amount of vesicles | CCV | DCV | NA | Total |
| --- | --- | --- | --- | --- | --- | --- |
| Adult hermaphrodite | 7 | Sum | 949 | 109 | 14 | 1072 |
|  |  | Percentage | 89% | 10% | 1% |  |
|  | 5 | Sum <sup>(1)</sup> | 837 | 85 | 11 | 933 |
|  |  | Percentage <sup>(1)</sup> | 90% | 9% | 1% |  |
| Dauer larvae | 8 | Sum | 948 | 148 | 12 | 1108 |
|  |  | Percentage | 86% | 13% | 1% |  |
|  | 5 | Sum <sup>(1)</sup> | 669 | 130 | 3 | 802 |
|  |  | Percentage <sup>(1)</sup> | 83% | 16% | 0% |  |

<sup>(1)</sup> analysis with a selected set of tomograms where only putative cholinergic NMJs with apparent dense projections and with > 100 vesicles were included.

### 4.2. Characterization based on image features

In order to perform a quantitative comparison of vesicles and for clarification about the ND vesicle class, we extracted a number of image features based on qualitative observations. Since DCVs appear larger, darker and further apart from the active zone compared to CCVs, we used radius (r), mean gray value (gv) and distance to active zone (distAZ) as primary features (Fig 2). In addition, the standard deviation of the gray value distribution of a vesicle (GVSD) is a measure for the contrast between membrane and core that is independent of the brightness of the vesicle.

**Fig 2.**
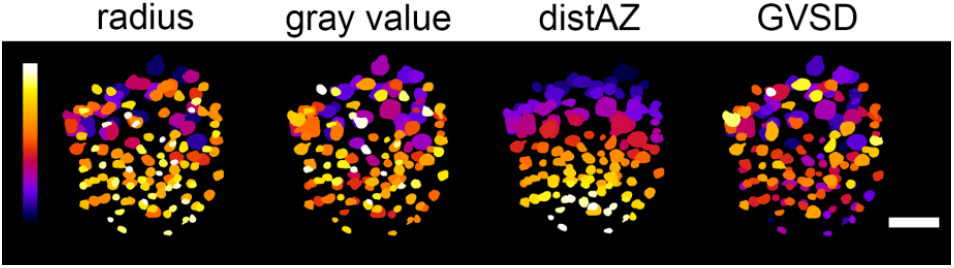
**Visualization of vesicle pool in a dauer larva tomogram with color coded image features:** inner radius, gray value, distance to active zone (distAZ), gray value standard deviation (GVSD). Clear core like characteristics are yellow-shifted, dense core like features are blue shifted. The active zone is located at the bottom.

Statistical analysis and comparison of ND vesicles with CCV and DCV suggests that synapses show no third type of vesicles and that NDs are in most cases CCVs (see S2 Fig). Hence, for further analysis, we proceeded with the assumption of two existing vesicle classes, CCVs and DCVs (Table 1). We tried to apply simple thresholding and manually weighted linear combinations of vesicle radius, distAZ and gv to separate DCVs from CCVs, but with limited success (see S2 Fig and S1 Text). Therefore, a machine learning approach was developed and applied, as described below. To generate training data, ND vesicles were first tried to be concordantly assigned to one group by two experts. Their label was only changed if they turned out to be DCV, as we programmed classifiers to treat the label ND as CCV. In difficult cases, a third expert was consulted. Still non-assignable vesicles (NA) were excluded from training by labelling them as error (see table 1: n = 12, ~ 1.1 % in dauer larvae; n = 14, ~1.3 % in young adult hermaphrodites).

### 4.3 Machine learning can distinguish vesicle types

To automatically classify vesicles using the previously calculated image features, we evaluated different machine-learning algorithms. We compared a support vector machine (SVM), random forest (RF), and k nearest neighbor (KNN) for their performance on differentiating between CCVs and DCVs using four image-based characteristics described above: radius (r), mean gray value (gv), standard deviation of the gray value (GVSD) and distance to the active zone (distAZ) (see Fig 3). 1994 manually labelled vesicles from 15 different tomograms (7 young adult hermaphrodite and 8 dauer larval tomograms) were used as training data. The features were extracted as in the first two steps of our classification macro (see Fig 3 and Methods section 6.3), saved as a table, and processed in a python script using the scikit-learn machine learning library [29]. Table 2 shows the results of the comparison of the different algorithms. Evaluation scores of the whole dataset were impractically high, as CCV greatly outnumber DCV. Therefore precision_DCV_, recall_DCV_ and F-score_DCV_ calculates scores for DCV specifically.

**Table 2.** Average performance of different machine learning classifiers

| classifier | accuracy | precisionDCV | recallDCV | F-scoreDCV |
| --- | --- | --- | --- | --- |
| SVM | $0.98 \pm 0.02$ | $0.95 \pm 0.07$ | $0.90 \pm 0.1$ | $0.92 \pm 0.07$ |
| Random forest<br>(t = 10) | $0.97 \pm 0.03$ | $0.88 \pm 0.12$ | $0.89 \pm 0.09$ | $0.88 \pm 0.08$ |
| Random forest<br>(t = 1500) | $0.97 \pm 0.02$ | $0.93 \pm 0.08$ | $0.86 \pm 0.1$ | $0.89 \pm 0.07$ |
| KNN<br>(k = 10) | $0.98 \pm 0.02$ | $0.92 \pm 0.09$ | $0.89 \pm 0.12$ | $0.90 \pm 0.09$ |

**Fig 3.**
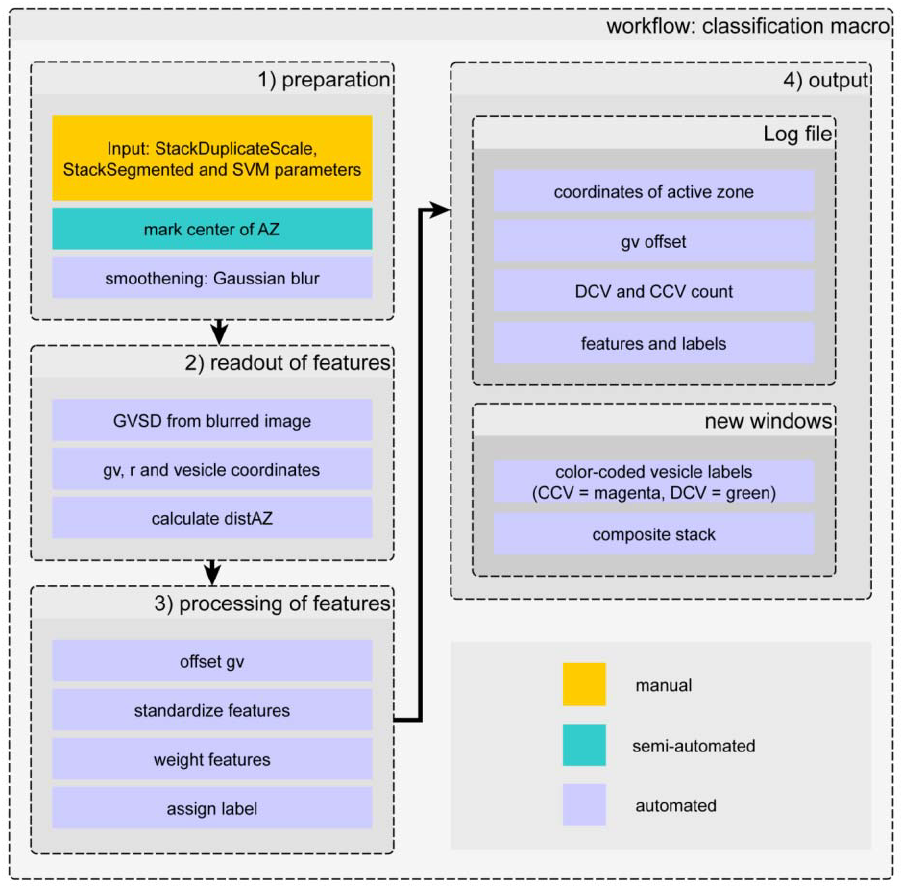
Workflow of the classification macro. 1) preparation: StackSegmented (registered vesicles) and StackDuplicateScale (scaled tomogram) from 3D ART VeSElecT [22] are used as input; the user is requested to mark the active zone via point-selection (semi-automated). 2) readout of features: gray value (gv), radius (r), vesicles coordinates and gray value standard deviation (GVSD) in the middle slice of vesicles are determined; the distance to active zone (distAZ) is calculated. 3) processing of features: offsetting, standardization and weighting of features followed by final assignment of labels. 4) output: includes a logfile and visualization of results as new image stack windows.

The comparison between SVM, random forest and KNN revealed that the SVM algorithm performs best on our classification problem. The SVM shows the lowest error rates in all four categories (see Methods): Accuracy, precision_DCV_, recall_DCV_ and F-score_DCV_. KNN shows second best results that were about 0-3 percent worse compared to the SVM. For the random forest variants, t = 1500 trees performed similarly or better than t = 10, except for recall_DCV_. Random forest (t = 1500) shows overall third best results with 3 percent worse results for recall_DCV_ and about 1 percent worse results for accuracy, precision_DCV_ and F-score_DCV_ compared to KNN. Resulting values of the SVM hyperplane delimiting DCVs from CCVs are shown in Table 3.

**Table 3.** SVM and standardization parameters transferred to the Fiji macro

| parameter | r | gv | distAZ | GVSD |
| --- | --- | --- | --- | --- |
| mean | 10.4 | 129.1 | 259.1 | 5.9 |
| SD | 3.1 | 4.7 | 118.5 | 1.7 |
| weight | -1.69 | 1.65 | -0.76 | 1.21 |

The final trained SVM classifier used the following weighting of the four features:

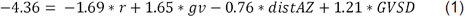

The weight for distAZ is smaller than for the other three parameters, which indicates that it is less important. Still, it contributes to the classifier, otherwise the resulting weight after training would be very small or zero. In such cases, it could be discarded from the training data.

In summary, the SVM algorithm in its linear form without kernelization outperformed all other classification methods including non-linear SVM and random forest (t = 1500) classification. Additionally, using a linear SVM has the advantage that the resulting classifier can easily be incorporated into a classification workflow by extracting the coefficients after successful training. For this purpose, a new vesicle classification macro in ImageJ/Fiji was developed to be consecutively applied after the modified 3D ART VeSElecT registration macro. The workflow of the classification macro is shown in Fig 3 and is described in detail in Materials and Methods.

### 4.4. Application to *C. elegans* dauer and adult hermaphrodite NMJs

We used the ground truth classification to quantify differences in the vesicle pool of *C. elegans* dauer and adult hermaphrodite NMJs in electron tomograms. Additionally, visual results of 3D ART VeSElecT and the classification macro for one exemplary tomogram of each analyzed developmental state are presented in Fig 4 to provide an intuitive impression of DCV and CCV distribution.

**Fig 4.**
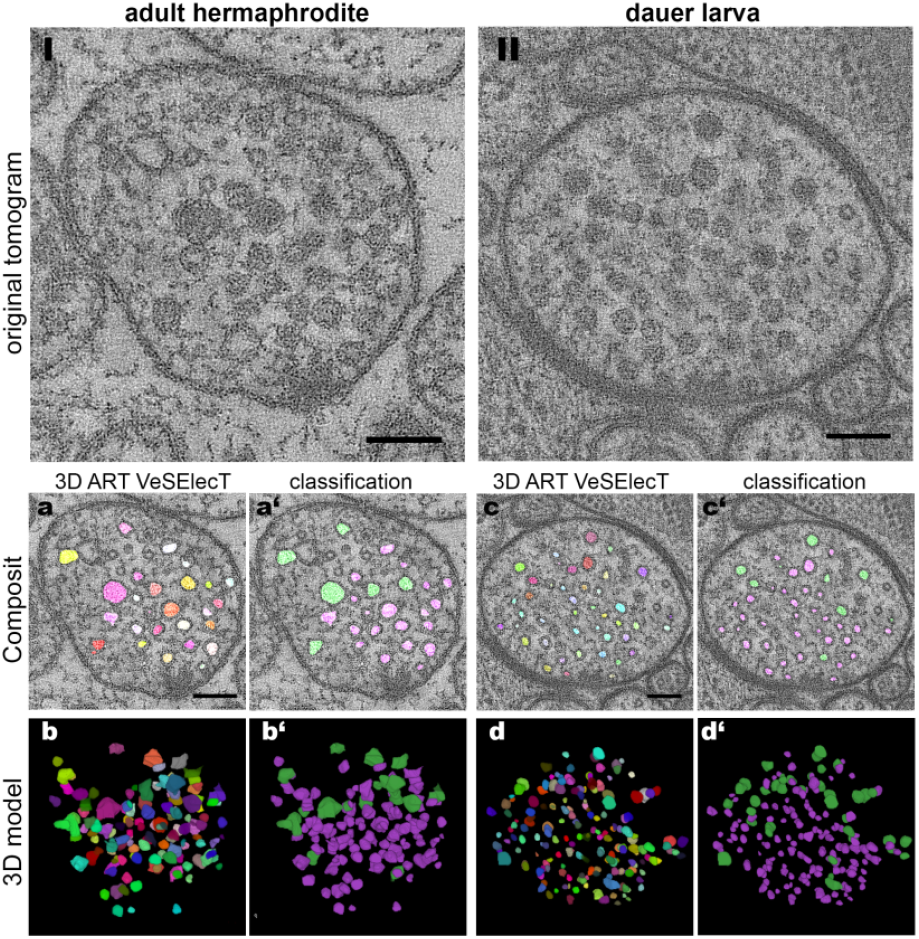
Comparison of the visual results from the adapted 3D ART VeSElecT and the consecutive classification macro. *Top:* I) and II) original tomograms of young adult hermaphrodite (I) and dauer larva (II). *Bottom:* a) and c) show *Composit* stacks of 3D ART VeSElecT (random colors), whereas a’) and c’) show the resulting *Composit* after classification of vesicles (CCV = magenta, DCV = green). b) and d) show the 3D model of the vesicle pool, b’) and d’) show the 3D model with color labels showing the assigned class. All scale bars = 100 nm.

The 3D models show that DCVs most often localize on the rim of the CCV cluster. CCVs themselves gather around the dense projections, electron dense structures attached to the cell membrane that indicate the active zone where synaptic vesicle fusion occurs. For further analysis of vesicle characteristics (radius, gray value, GVSD and distAZ), results are shown as violin plots in Fig 5.

**Fig 5.**
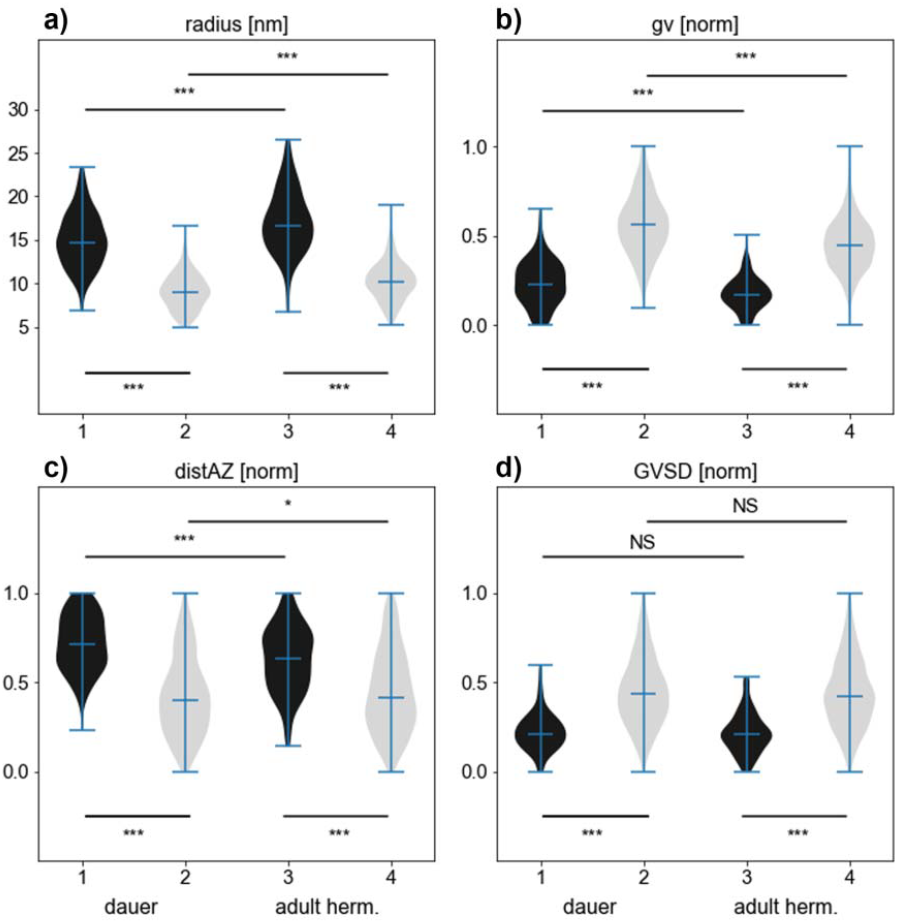
Comparison of DCVs (black) characteristics with CCVs (light gray) characteristics. Violin plots show a) inner vesicle radius in nm, b) gray value (gv), c) distance to active zone (distAZ) and d) gray value standard deviation in central slice of each vesicle (GVSD). Panels b)-d) show normalized values of n_h_ = 7 tomograms of N2 young adult hermaphrodites; n_d_ = 8 tomograms of N2 dauer larvae. All plots are based on expert curated manual ground truth classification.

Fig 5a shows highly significant differences in CCV and DCV size in both developmental states. In the following, the radius always refers to our original measured results, which is the “inner radius” of all vesicles, as described in [22]. To make vesicle size comparable to other published results, the measured radius must be multiplied by two to receive the inner diameter and increased by 9 nm to include vesicle membrane thickness [22]. This estimated diameter is only an approximation and cannot be directly compared to manually measured data due to the different way how both are measured [22].

CCVs of dauer larvae show a median inner radius of 9.0 nm, whereas DCVs have a median inner radius of 14.7 nm, which results in a significant difference of 5.7 nm. However young adult hermaphrodites’ median inner radius of DCVs is 16.7 nm, hence they have a ~ 5.4 nm bigger radius than CCVs with a median inner radius of 10.3 nm. Noticeably, both vesicle types are slightly but significantly bigger in young adult hermaphrodites compared to dauer larvae. The radius of hermaphrodite CCVs is 1.3 nm larger compared to dauer larvae, whereas the radius of hermaphrodite DCVs is 2 nm larger compared to dauer larvae.

In all cases, DCVs have significantly darker normalized gray value intensities in comparison to CCVs, as shown in Fig 5b. Young adult hermaphrodites show a normalized gv of 0.17 for CCVs and 0.45 for DCVs, whereas dauer larvae have a normalized gv of 0.23 for CCVs and 0.56 for DCVs. Hence, vesicles of dauer larvae tomograms possess significantly darker vesicles (both CCVs and DCVs) in comparison to hermaphrodite tomograms. In contrast, GVSD (gray value standard deviation in the central slice of each vesicle) does not differ significantly between developmental states, but shows highly significant differences of ~ 0.22 in GVSD values of CCV to DCV for both developmental states (Fig 6d).

**Fig 6.**
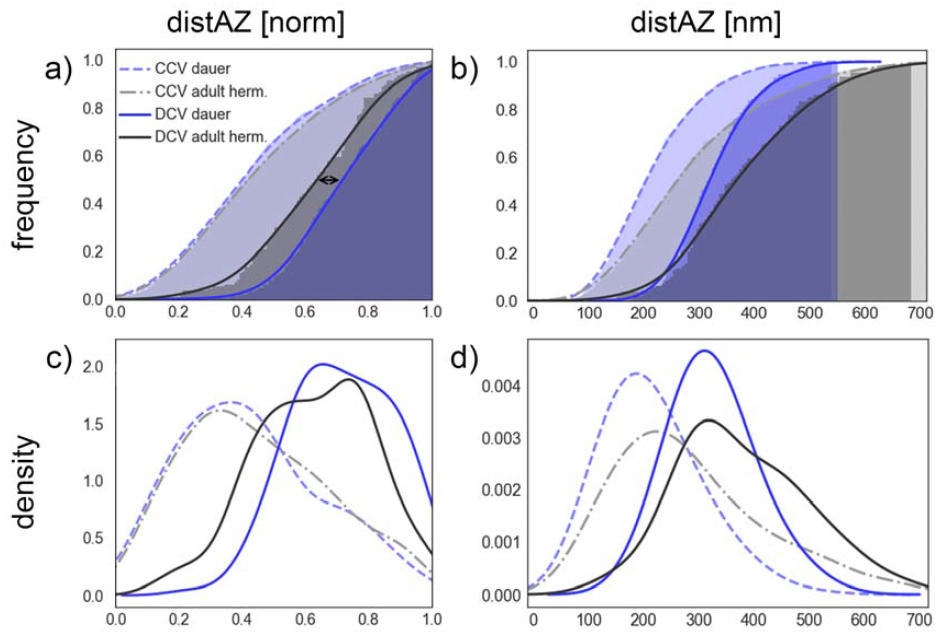
Cumulative histogram (a, b) and kernel density estimation plots (c, d; bandwidth = 0.07*maximum value of total dataset) of distances to the active zone (distAZ). Given are normalized distances (left) and real values in nm (right). CCVs are shown as dashed line and DCVs as continuous line. Adult hermaphrodite vesicles are presented in shades of gray, whereas dauer larval vesicles are shown in violet. All plots are based on expert curated manual ground truth classification.

The normalized distance of CCVs to the active zone distAZ is smaller (0.42 for young adult hermaphrodites and 0.4 for dauer larvae) than that of DCVs (young adult hermaphrodites: 0.64, dauer larvae: 0.72) in both larval stages (Fig 6c). The normalized distances of DCVs to the AZ, of young adult hermaphrodites in comparison to dauer larvae, show a highly significant difference of 0.08 (normalized). For better visualization of this observation, distances are visualized as histogram plots in Fig 6.

Fig 6a and c illustrate that normalized distances of CCVs of both developmental states are almost identical, whereas distances of DCVs differ significantly in both plots. DCVs show a systematic discrepancy in their localization: DCVs in young adult hermaphrodites cluster closer to the active zone than DCVs of dauer larvae. Histograms in Fig 6b and d shows results of distAZ in real distance measured in nm. Interestingly, this statistic shows that vesicle pools in dauer larvae have a generally smaller maximal distAZ in young adult hermaphrodites. The largest distance of dauer larvae vesicles from the active zone is about 550 nm whereas the largest distance of vesicles in young adult hermaphrodites is about 700 nm. It should be noted that distAZ as derived from tomograms should be considered as an estimate of the actual distance of vesicles to the active zone, since a single tomographic slice may not contain the full information on the actual position and shape of the active zone.

### 4.5. Error rates and cross validation

To generate the ground truth data, a consensus label was created (N = 15 tomograms, 8 dauer larva and 7 hermaphrodite tomograms): First, labels were assigned independently by two biologists. Differing labels were re-examined. In case of no agreement, a third examiner was employed. NA vesicles (~1 %) were excluded from SVM training. The initial assignment by individuals was compared with this consensus label to determine the manual error rate (Table 4).

**Table 4.** manual vs classification macro (SVM performance)

|  | presicion<sub>DCV</sub> | recall<sub>DCV</sub> | F-score<sub>DCV</sub> |
| --- | --- | --- | --- |
| Manual <sup>(1)</sup> | 0.98 ± 0.05 | 0.96 ± 0.08 | 0.97 ± 0.05 |
| SVM <sup>(2)</sup> | 0.95 ± 0.07 | 0.90 ± 0.10 | 0.92 ± 0.07 |

- (1) Compares two independent manual assignments to the consensus label (ground truth) - (2) Compares the SVM assignment to the ground truth

Manual annotation shows higher scores than the SVM. Manual assignment of vesicle types shows 98 % precision_DCV_, whereas the SVM reaches 95 % precision_DCV_. The results for recall_DCV_ is 96 % for manual assignment and 90 % for the SVM. The F-score is the harmonic mean of precision and recall (see Materials and Methods) and shows a mean result for the error rate of 97% for manual and 92% for automated classification. Results for precision and recall are additionally shown as scatterplot in Fig 7.

**Fig 7.**
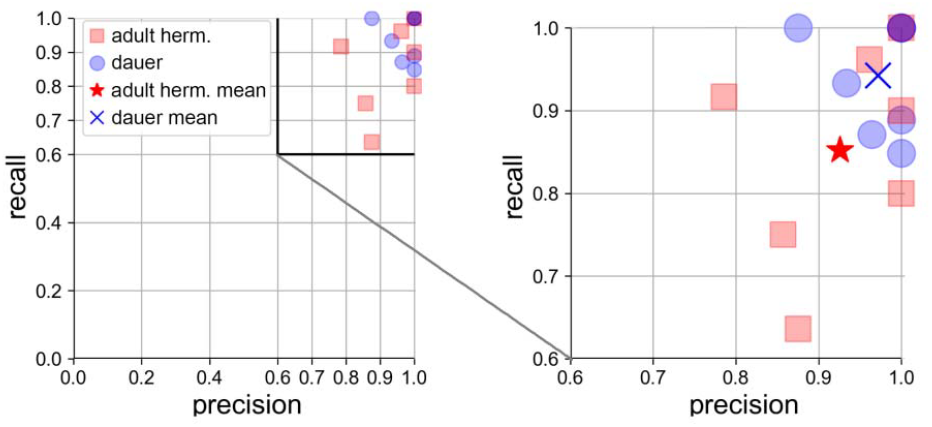
SVM error rates as precision_DCV_ and recall_DCV_ plot. Each data point represents one validation tomogram. Red squares: adult hermaphrodite, blue circles: dauer larvae. Right: magnified detail plot. Red star indicates mean of hermaphrodite data: precision_DCV_ = 0.93, recall_DCV_ = 0.85. Blue cross indicates mean of dauer data: precision_DCV_ = 0.97, recall_DCV_ = 0.94. 3 dauer tomograms and 1 hermaphrodite tomogram were classified without error.

The SVM classifier was cross-validated against each tomogram using all other tomograms as training data. Fig 7 shows error rates of cross validation of automated SVM classification. Hermaphrodite tomograms are represented as red squares and dauer larvae tomograms are represented as blue circles. Mean classification results are shown as red star for young adult hermaphrodites (precision_DCV_ = 0.93, recall_DCV_ = 0.85) and as blue cross for dauer larvae tomograms (precision_DCV_ = 0.97, recall_DCV_ = 0.94). Dauer larvae tomograms have in average better results than hermaphrodites and cluster in the right upper corner. 3 dauer tomograms and one young adult hermaphrodite tomogram were classified without any error and overlay in the right upper corner at precision_DCV_ = 1.0, recall_DCV_ = 1.0.

## 5. Discussion

Automated methods for image analysis are of great importance to reduce manual labor in the long term and prevent biased analysis of data, hence making double blind analysis unnecessary. This is confirmed by our own experience, where one re-analysis of the same tomogram by the same expert revealed one differential interpretation of CCV and DCV and 6 other non-matching assignments on n = 134 vesicles in total. Such deviations in manual analysis may have a great influence on the outcome. For this reason, we had two experts manually assign all vesicles independently. In case of inconsistency, both persons came to a mutual agreement or consulted a third expert in the decision process. Still unassignable vesicles were excluded from analysis. Since this evaluation process is very elaborate and time-consuming, a long-term automated solution is inevitable. As far as we know there is no other approach for the specific task of automated vesicle classification in electron tomograms. We present here a new methodological approach that is applied subsequent to a modified version of our previously published Fiji macro 3D ART VeSElecT [22]. First, a Fiji macro was written to read out important characteristics. Those were, together with manual labelling, used to train machine learning classifiers in python. Extracted machine learning parameters of an SVM completed the new Fiji macro to classify vesicles and visualize classes through 3D color labelling. Our tool can thus be used for automated vesicle classification. We used our ground truth label to compare two different developmental states of *C. elegans*, young adult hermaphrodites and dauer larvae.

Manual analysis revealed a hypothetical third type of vesicles, called non-determinable vesicles (ND) which are most likely CCVs with no perfectly clear inner core. This seems reasonable, since they resemble CCV characteristics in radius and distance to the active zone and only differ significantly in their gray value, which lies among gv intensities of DCVs and CCVs (S2 Fig). This observation can be explained by either biological or technical reasons. On the one hand, medium-gray vesicles could be DCVs with partially released cargo, e.g. after kiss-and-run [30–32]. Overall this does not explain why NDs have the same size as CCVs, since DCVs are known to be considerably larger [23]. On the other hand, vesicles may appear darker in electron micrographs because of lower quality of tomograms at some locations. Lacking precision of the tomogram overlay of the two single tilted tomograms to receive one double tilted tomogram could result in darker, more blurry vesicles with no perfectly clear core.

The comparison of DCV number in dauer larvae and young adult hermaphrodites revealed differences in the ratio of DCVs (9 % in young adult hermaphrodites and 16 % in dauer larvae), when comparing only cholinergic synapses with apparent dense projections and at least 100 vesicles in the NMJ. Furthermore we found that DCV ratio correlates with cell size change throughout the tomogram (S4 Fig). A large cell size change indicates inclusion of decentral regions of the synaptic bouton in the tomogram. We assume that inclusion of more remote areas in the tomogram is more likely in dauer larvae, since the overall size of the synaptic bouton is smaller (mean diameter ± SD of cross section: 452 ± 85 nm) compared to young adult hermaphrodites (mean diameter ± SD of cross section: 542 ± 97 nm). Therefore, standard sized 250 nm sections, typically prepared for electron tomography, naturally have a higher chance of catching decentral regions in dauer larvae NMJs. For a more comprehensive understanding, serial section tomography for visualization of complete synaptic boutons would be necessary.

Additionally, we found that young adult hermaphrodites show significantly larger vesicles (CCVs and DCVs) and diameters of synapses in comparison to dauer larvae. This observation could either be a biological phenomenon of dauer larvae or a side effect of the sample preparation. Dauer larvae have a distinct cuticle that shields the animal from adverse environmental conditions, like heat and drought. This could lead to adverse effects during fixation (freeze substitution) of the sample, even though we did not observe any sign of deficient sample fixation in dauer larvae. Divergent synaptic vesicle sizes have already been found in NMJs of other organisms, like rats [33] and zebrafish [23]. For zebrafish a dependency of vesicle size on the maturation stage of NMJs was assumed that might be related to the functional transition of these synapses to control different swimming behavior [23]. Differences in synaptic vesicle sizes during normal development have not been observed for *C. elegans*, to our knowledge. However, divergent vesicle sizes could also be related to different moving behavior of dauer larvae that are well known for their unique nictation behavior [9]. Since manual measurements are often inaccurate, smaller deviations in vesicle sizes are only observable in extensive analysis, therefore automated approaches are inevitable. Our previously published macro 3D ART VeSElecT proved to be successful for distinguishing vesicle size differences [22]. This approach in combination with our here presented extended version for automated classification may be the key to reveal so far undistinguishable differences in the future.

Interestingly, although nervous cells in dauer larvae are typically smaller, segregation of DCVs and CCVs is more distinct. Our interpretation is that DCVs in dauer larvae locate further away from the active zone on the rim of the synaptic vesicle pool compared to young adult hermaphrodites. Differences in this characteristic of synaptic architecture could be explained by a shift towards a higher ratio of DCVs in a non-fusion competent resting state in dauer compared to young adult hermaphrodites. Moreover, the location of DCVs closer to the cell margin in dauer compared to young adult hermaphrodites could indicate the need for faster peptide release, and their increased number could compensate for lower volume.

For automated classification of vesicles, different machine learning algorithms were tested. The SVM showed best results in comparison to random forest or k-nearest neighbor classification. Furthermore, the SVM works regardless of dimensionality, can classify non-linearly separable classes by using the kernel trick, enables an easy integration of the trained linear classifier into the macro and generally can be used for assignment into more than two classes [34].

Cross-validation showed very good results for automated classification of vesicles. Interestingly, dauer larval tomograms showed better results than young adult hermaphrodite tomograms, which could be related to the differing gray value intensities of the tomograms. Hence high contrast vesicles were easier to automatically recognize and showed more distinct results for GVSD and gv. Another reason could be the more pronounced positional segregation of the two vesicle types for dauer larvae.

We suggest that our pre-trained SVM can be used for DCV and CCV classification in other, similar tomograms, and provide the Fiji classification macro including the trained parameters. In addition, users can train a SVM on their own data using the python script we provide, and integrate the parameters into the Fiji macro, which is quickly achievable via an input dialog.

Our ImageJ classification macro and the python script, for retraining of the classifier, a tutorial and test datasets are available at: https://www.biozentrum.uni-wuerzburg.de/bioinfo/computing/3dart-veselect/.

## 6. Conclusion

We developed a new automated approach to classify vesicles and to quantify their properties from electron tomograms. We combined machine learning with an extension of our previously developed vesicle segmentation workflow 3D ART VeSElecT to reliably distinguish clear core from dense core vesicles using image-based features. We apply this method to electron tomograms of *C. elegans* NMJs in young adult hermaphrodites and dauer larvae. Using our ground truth data, we find an increased fraction of dense core vesicles (~ 16 % vs ~ 9 %) in tomograms matching certain characteristics (cholinergic synapses with > 100 vesicles of a central slice through the AZ area), as well as significantly reduced vesicle size and a higher distance of dense core vesicles to the active zone in dauer larvae compared to young adult hermaphrodites. Our approach is not limited to *C. elegans* and can be easily adapted to study differences in synaptic and other vesicles in electron tomograms in various settings and many other model organisms, e.g. in *Danio rerio* [22].

## 7. Material and Methods

### 7.1 Electron Microscopy

#### Animals

C. elegans wild-type (Bristol N2) were used and maintained using standard methods [4]. For the experiments, young adult hermaphrodite worms as well as dauer larvae were used.

#### Purification of dauer larvae

For segregation of dauer larvae from other states of the C. elegans life cycle, treatment with sodium dodecyl sulfate (SDS) buffer was used [35]. Firstly, worms were washed from the agar plates using M9 medium [36]. The worm solution was then transferred into 50 ml tubes and centrifuged for 5 min at 2000g at 4°C. Afterwards, the supernatant was removed and the pellet was dissolved in 50 ml 1 % SDS solution and incubated on a shaker for 15 min. SDS was then removed by centrifuging (2000g, 5 min, 4°C) and subsequent discarding of the supernatant while retaining 5 ml of solution. The tube was then refilled with distilled water and the procedure repeated 2-3 times. For separation of dauer larvae from dead material, the solution was pipetted into a pit on an unseeded agar plate. Once the solution had dried, living dauer larvae were able to crawl away from the pit. After 1-3 hours, emigrated worms were washed off with M9 medium.

#### High-pressure freezing, freeze-substitution and electron tomography

HPF, FS and electron tomography were performed as previously described [22]. Tilt series were recorded from at least +65° to −65° but not exceeding +70° to −70°.

### 7.2 Improvements to 3D ART VeSElecT

We extended our tool 3D ART VeSElecT to calculate various image features of detected vesicles to be used for classification. The workflow is shown in Supplementary Figure S1, with changes compared to the previous version highlighted by a black box. The overall workflow of 3D ART VeSElecT remained as previously published [22]. One change in the preprocessing step is the determination of the mean radius instead of volume, since the radius or diameter is typically used when comparing vesicle size. Furthermore, automated calculation of the cell volume was added. Since semi-automated selection of the cell outline by the user was already included in the macro, this selection is now basis for calculating the analyzed volume in 3D (which corresponds to cell volume if the selection is taken along the cell membrane, as described in the 3D ART VeSElecT tutorial).

### 7.3 Classification Macro

The workflow of the new Fiji macro for classification of DCV and CCV in electron tomograms of synapses comprises four main steps: 1) preparation, 2) reading out features, 3) processing of features followed by labelling of vesicles, and 4) output. In the preparation step, StackDuplicateScale and StackSegmented, which result from the 3D ART VeSElecT recognition macro, are read into the classification macro. StackDuplicateScale is the scaled original tomogram, StackSegmented contains the registered vesicles. First, the macro opens a preparation window for changing the names of read-in stacks (StackDuplicateScale and StackSegmented) and setting parameters for standardization and adaption of SVM weights. In case of adaptation of the SVM algorithm to another user’s needs, values can easily be changed through the input window. In a second step, four characterizing vesicle features are computed from the images: radius, gv, GVSD and distAZ. GVSD is a value for standard deviation of gray values in the central slice of each vesicle, since CCVs have a higher gv variance than DCV. In the third step, features are processed, which includes applying an offset to gv (if the darkest vesicle is lighter than a threshold), standardization of parameters, weighting features based on SVM weights, and finally the assignment of labels according to the identified class (CCV = magenta, DCV = green). In the last step, the output is created, including a logfile and visual output shown in new windows. The logfile includes all measured characteristics (e.g. features, classification labels (CCV = 1, DCV = 0), number of vesicles in each class). The visual output shows, besides the input stacks (“StackSegmented” and “StackDuplicateScale”), an additional image stack, named “label” which corresponds to the StackSegmented but displays color-coded vesicles according to their classification. Furthermore a Composit stack, named “CompositLabel” (overlay of original tomogram and the “label” stack) is created.

### 7.4 Comparison of different types of machine learning classifiers

We compared a support vector machine (SVM), random forest (RF, using t = 10, t = 1500; t indicates the number of trees) and k nearest neighbor (KNN, k = 10 neighbors) for their performance on differentiating between CCVs and DCVs based on four image-based characteristics: radius (r), mean gray value (gv), standard deviation of the gray value (GVSD) and distance to the active zone (distAZ). Random forest uses decision trees to classify samples: Comparable to a flowchart, samples are divided based on their features. A random forest utilizes the majority classification of t decision trees each grown on randomly selected training data and split using the best randomly selected feature. Generalization error is lower than for individual decision trees. Performance, but also runtime increases with t. KNN classifies samples based on the majority of their k nearest training samples in feature space. Linear SVM (C = 1) showed the best average classification results (see Table 2). C determines the penalty for falsely classified training data points and therefore indicates bias-variance trade-off, since bias decreases with C [37]. The SVM will be described more accurately below. Parameters of the SVM (Table 3) were determined using all input tomograms as training data and transferred to a Fiji-Macro (Figure 3) to enable classification downstream of the 3D ART VeSElecT registration macro.

### 7.5 Support Vector Machine (SVM)

A linear support-vector machine (SVM) divides two classes by a hyperplane. As such, the SVM is a binary classifier dealing with linear classification problems. A one-versus-all approach and kernelization can extend classification to non-linear problems and into more than two classes, but will not be discussed here since the classification power of the SVM is sufficiently strong, see results. With the aim of minimizing generalization error, the optimization objective of the SVM is to maximize the margin between the hyperplane and the closest data points of the two classes. Those eponymous support-vectors lie on the positive and negative hyperplane, which are equidistant to the separating hyperplane [37]. As the linear SVM is parametric, the equation of the separating hyperplane can be transferred independently of the training-data (see Results, Table 3 and Equation 1), with x_i_ being the features and ωi the assigned feature-weights. All data points with 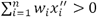 are assigned to class 1 (CCV), whereas all data points with 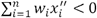 are assigned to class −1 (DCV).

### 7.6 Implementation of machine learning algorithms

A python 3 script (S3 Fig) reads in the CSV files containing the training datasets and applies classifiers from the scikit-learn library [29]. The manual labels are transformed into a feature array: Particles marked “D” (DCV) are relabeled “−1”. Particles marked “E” (error) are removed. All other markings are relabeled “1”, including “C” (CCV) and “N” (not determinable), since non-determinable particles show more similarities to CCV (S2 Fig).

Classifiers are evaluated by cross-validation: Data from one CSV file are used as validation data, all other files are concatenated and used as training data (there are as many training-validation combinations as input CSV files). For each combination, features are standardized using mean and standard deviation (sd) of the training data and passed to the classifiers, namely linear SVM, random forest and KNN. An evaluation function determines accuracy of all vesicles and precision, recall and F-score of DCV only (see equations (2) - (4)). Evaluation results are returned as pandas.DataFrame [38] and as CSV file.

**Fig 8.**
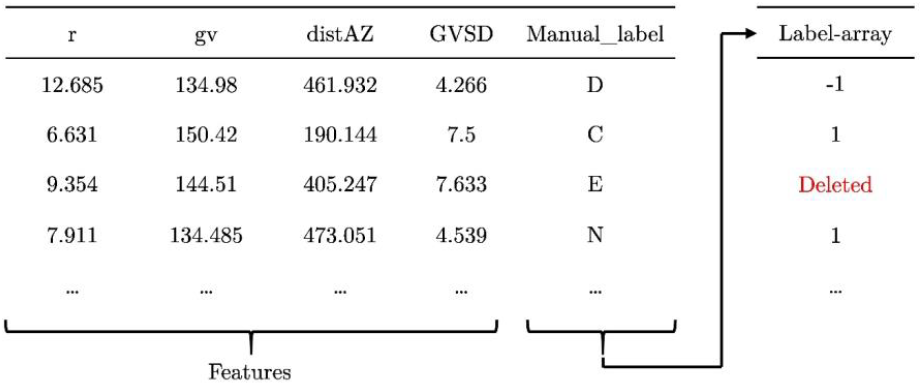
Format of input CSV file. First four columns are the four features, the fifth column is the manual labels. The python script rewrites manual labels into a binary label-array: “D” (DCV) is rewritten to −1, “E” (error) is removed, and everything else is rewritten as “1” (CCV).

Equations:

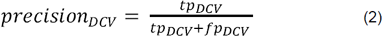

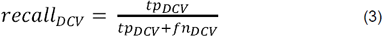

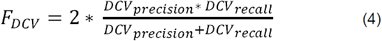

tp: true positive; fp: false positive; fn: false negative.

The runtime of the training script for the different classifiers was 5 s for t=10 trees and 140 s for t=1500 trees. The ImageJ classification macro takes 33s for the calculation, plus the time required to manually click on the active zone. All computations were performed on a 16-core AMD Opteron 1.7 GHz server with 128 GB RAM running Ubuntu Linux 16.04. In comparison, manual classification of vesicles takes an experienced user about 1 h per tomogram. The time required to set up the python environment and prepare the training data depends on the level of experience of the user, but overall there should still be at least a factor of 10 (one order of magnitude) of speed-up with reasonably large data sets of a few tomograms, as in our case. In addition, automated workflows as reported here increase reproducibility compared to manual assignment.

